# Autographa californica Multiple Nucleopolyhedrovirus *orf13* is Required for Efficient Nuclear Egress of Nucleocapsids

**DOI:** 10.1101/2020.07.13.201756

**Authors:** Xingang Chen, Xiaoqin Yang, Chengfeng Lei, Fujun Qin, Jia Hu, Xiulian Sun

## Abstract

Autographa californica multiple nucleopolyhedrovirus (AcMNPV) *orf13* (*ac13*) is a conserved gene in all sequenced alphabaculoviruses. However, its function in the viral life cycle remains unknown. In this study we found that *ac13* was a late gene and that the encoded protein, bearing a putative nuclear localization signal motif in the DUF3627 domain, colocalized with the nuclear membrane. Deletion of *ac13* did not affect viral DNA replication, gene transcription, nucleocapsid assembly or occlusion body (OB) formation, but reduced virion budding from infected cells by approximately 400-fold compared with the wild-type virus. Deletion of *ac13* substantially impaired the egress of nucleocapsids from the nucleus to the cytoplasm, while the number of occlusion-derived viruses embedded within OBs was unaffected. Taken together, our results indicated that *ac13* was required for efficient nuclear egress of nucleocapsids during virion budding, but was dispensable for OB formation.

**IMPORTANCE:** Egress of baculovirus nucleocapsids from the nucleus is an essential process for morphogenesis of mature budded viruses, which is required to spread infection within susceptible cells and tissues. Although many viral and host proteins are required for nucleocapsid egress, the specific mechanisms underlying this process in baculoviruses remain somewhat enigmatic. In the present study, we found that the *ac13* gene, in addition to *ac11, ac51, ac66, ac75, ac78, gp41, ac93, p48, exon0* and *ac142*, was required for efficient nuclear egress of nucleocapsids. Our results contribute to a better understanding of nucleocapsid egress in baculoviruses.

## INTRODUCTION

The Baculoviridae are a large family of insect-specific viruses with circular, covalently closed, double-stranded DNA genomes 80–180 kb in size and encoding 89 to 183 genes (1, 2). Based on their genome sequences, baculoviruses can be divided into four genera: *Alphabaculovirus, Betabaculovirus, Gammabaculovirus*, and *Deltabaculovirus* (3). Alphabaculoviruses can be further subdivided into Group I and Group II viruses (4). The most notable differences between these two groups are that Group I nucleopolyhedroviruses (NPVs) use GP64 as their budded virus (BV) fusion protein, whereas Group II NPVs lack *gp64* and use the F protein (5). Autographa californica multiple nucleopolyhedrovirus (AcMNPV) is the archetype species of *Alphabaculovirus*.

Baculovirus infection produces two distinct viral phenotypes: BVs and occlusion-derived viruses (ODVs) (2). BVs are responsible for spreading infection within susceptible insect cells and tissues, whereas ODVs initiate primary infection in the midgut epithelia of infected insects and are transmitted among insects (6). Transcription and replication of viral DNA and assembly of nucleocapsids occur in a structure called the virogenic stroma (VS) (7, 8). Synthesized nucleocapsids are transported from the VS to the ring zone, and then egress from the nucleus to cytoplasm and bud from the plasma membrane to form BVs. Subsequently, nucleocapsids retained in the ring zone of the nucleus are enveloped by intranuclear microvesicles to form ODVs, which are then embedded within the polyhedrin to form OBs (2).

Most DNA viruses, including herpesviruses and baculoviruses, replicate and assemble their nucleocapsids in the nucleus (2, 9). Egress of nucleocapsids is indispensable for formation of mature virions and viral pathogenicity. This process also represents a good target for disrupting viral infection. The mechanism through which herpesvirus nucleocapsids egress has been well characterized (9, 10). By contrast, the mechanism of baculovirus nucleocapsid egress remains unclear. According to previous reports, host proteins including the actin cytoskeleton, N-ethylmaleimide-sensitive fusions proteins and endosomal sorting complex required for transport-III (11-13) as well as viral proteins including Ac11, Ac51, Ac66, Ac75, Ac78, GP41, Ac93, P48, EXON0 and Ac142 are required for nucleocapsid egress (14-25). Deletion of *ac11, ac75, ac78, gp41, ac93, p48*, or *ac142* abrogated egress of nucleocapsids from the nucleus. By contrast, loss of *ac51, ac66* or *exon0* reduced the efficiency of nucleocapsid egress. According to previous reports, the nucleocapsids of BVs were ubiquitinated at much higher levels than those of ODVs, indicating that nucleocapsid ubiquitination (potentially catalyzed by the viral E3 ubiquitin ligase EXON0) may play a key role in nucleocapsid egress (26). Exploring genes associated with nucleocapsid egress is important to elucidate the mechanism of nucleocapsid egress in baculoviruses.

*ac13*, encoding a protein of 327 amino acids with a putative molecular mass of 38.7 kDa (9), is a conserved gene in all sequenced alphabaculoviruses. However, the function of *ac13* in the viral life cycle remains unknown. To date, only a few studies have examined *ac13* and its orthologs. Transcriptomic sequencing showed that *ac13* was regulated by an early promoter and a late promoter (27). InterProScan (28) and NCBI Conserved Domain Search (29) analyses revealed that Ac13 contained a DUF3627 protein domain of unknown function, which was conserved in all alphabaculovirus but not betabaculovirus orthologs. *bm5*, a homolog of *ac13* in Bombyx mori NPV (BmNPV), was seemingly nonessential because the viral life cycle appeared normal when it was deleted (30, 31). However, a recent study showed that although deletion of *bm5* did not affect viral DNA replication, it decreased BV and OB production (32).

In the present study, we investigated the function of *ac13* in the baculovirus life cycle. First, temporal analysis of transcription and transcription initiation sites (TSSs) showed that *ac13*, with an early and a late promoter, was transcribed during both the early and late phases of infection. However, the Ac13 protein was only detected during late infection and colocalized with the nuclear membrane. In addition, we determined the roles of *ac13* in BV production, viral DNA replication, viral gene transcription and OB morphogenesis. Our results indicated that *ac13* was not essential for viral DNA replication, gene transcription, nucleocapsid assembly or OB formation. However, its absence reduced the efficiency of nucleocapsid egress and decreased the production of BVs.

## RESULTS

### 1. *ac13* is a late viral gene

Transcriptomic analysis showed that two different TSSs were located upstream of the *ac13* translation initiation codon (27). Temporal transcription patterns showed that the product of *ac13* was detected as early as 6 h post infection (h p.i.) and persisted up to 48 h p.i. (Fig. 1A). Rapid amplification of 5’ cDNA ends (5’ RACE) revealed that the TSSs mapped to the first G of the atypical baculovirus early promoter motif GCAGT, located 217 nt upstream of the *ac13* open reading frame (ORF) start codon, and the first A of the typical late promoter motif TAAG, located 56 nt upstream of the *ac13* ORF start codon (Fig. 1B). These results indicated that expression of *ac13* was regulated by an early and a late promoter, and that the gene was transcribed during early and late infection of host cells.

**FIG 1.**
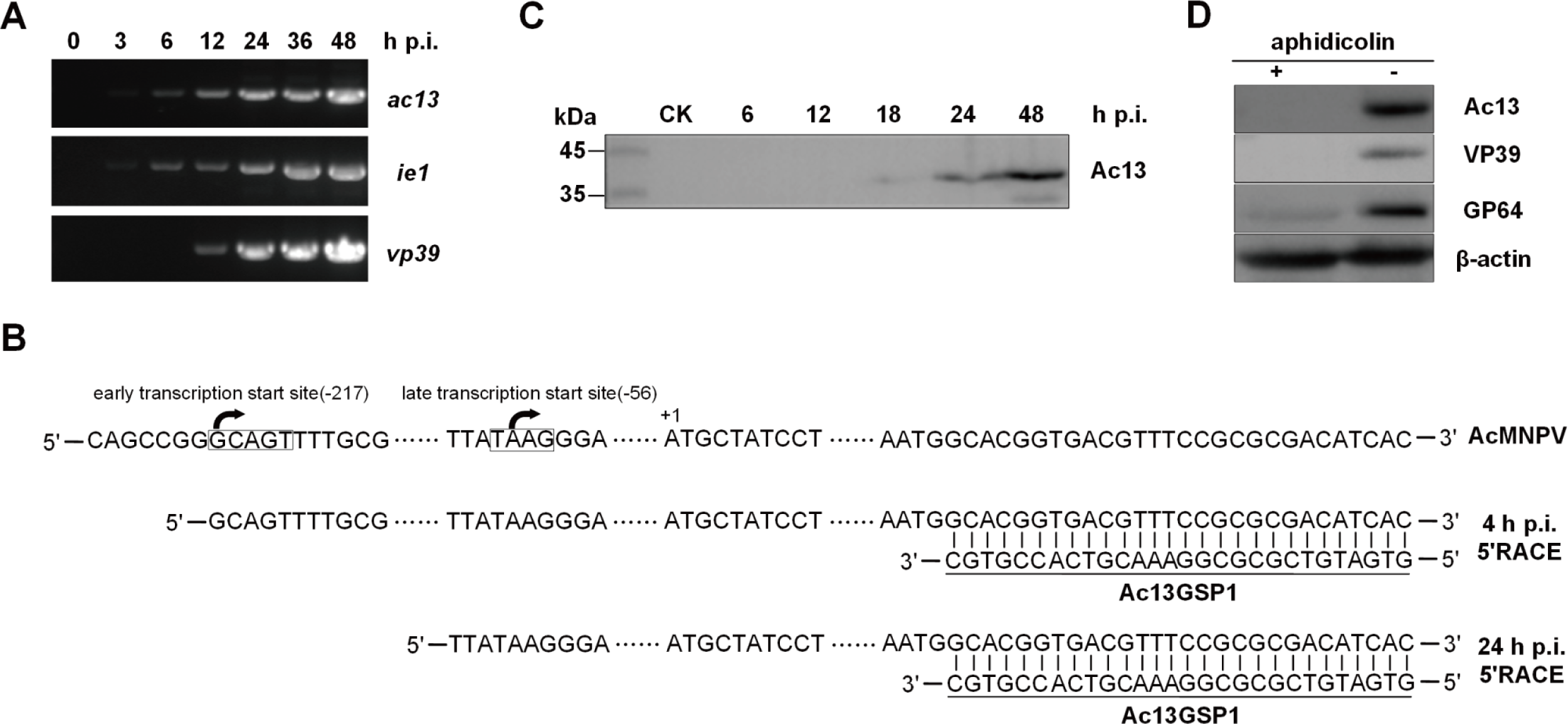
Transcription and expression analysis of *ac13* in Sf9 cells. (A) RT-PCR analysis of *ac13* transcription. Total RNA was extracted from AcMNPV-infected Sf9 cells at the indicated time points and were amplified the transcripts of *ac13, ie1* and *vp39*, respectively. (B) 5’ RACE analysis of *ac13* TSS. Total RNAs were extracted from AcMNPV-infected cells at 4 and 24 h p.i. and subjected to 5’ RACE analysis. The late promoter (TAAG) and the early promoter (GCAGT) were denoted in box and the two TSS were shown with arrowhead. (C) Western blots analysis the temporal expression of Ac13. The Sf9 cells, infected with AcMNPV at an MOI of 10, were harvested at the indicated time points and detected with anti-Ac13 antibody. (D) Western blots analysis of the expression of Ac13 with aphidicolin. The AcMNPV-infected cells were treated with 5 μg/ml aphidicolin (+) or DMSO control (-) at 0 h p.i. and then processed at 24 h p.i. and detected with anti-Ac13. The anti-VP39, anti-GP64 and anti-actin were used as controls.

The temporal expression profile of Ac13 was determined by western blotting of AcMNPV-infected cells at designated time points. A band of approximately 39 kDa, close to the predicted molecular mass of Ac13, was detected from 18 to 48 h p.i. (Fig. 1C). To further determine whether Ac13 was expressed during late infection, Ac13 was detected in AcMNPV-infected cells via the presence of aphidicolin, which inhibits viral DNA replication and thus prevents viral late gene expression. No Ac13 expression was observed in aphidicolin-treated cells, while Ac13 was detected in control cells treated with dimethyl sulfoxide (DMSO) (Fig. 1D). Expression of VP39 was only detected in DMSO-treated cells, while expression of GP64 was detected both in aphidicolin-treated and DMSO-treated cells (Fig. 1D). Together these results showed that despite transcription of *ac13* during early and late infection, it was a late viral gene.

### 2. Ac13 is predominantly localized to the nuclear membrane

To investigate the function of Ac13 in the viral life cycle, its subcellular localization was analyzed by confocal microscopy. Sf9 cells infected with vAc^*ac13Flag*REP^-*ph* at a multiplicity of infection (MOI) of 5 were fixed at 12, 18, 24, 36, 48 and 72 h p.i. Ac13 was detected by immunofluorescence using confocal microscopy. As shown in Fig. 2A, Ac13 fluorescence was predominantly localized to the nuclear rim and mainly colocalized with the nuclear lamina of the nuclear membrane from 12 until 72 h p.i.. Immunoelectron microscopy was used to assess the location of Ac13 in cells infected with vAc^*ac13Flag*REP^-*ph*. At 48 h p.i., colloidal-gold-labeled Ac13 was predominantly localized to the perinuclear and nuclear membranes (Fig. 2B). These results agreed with the immunofluorescence data.

**FIG 2.**
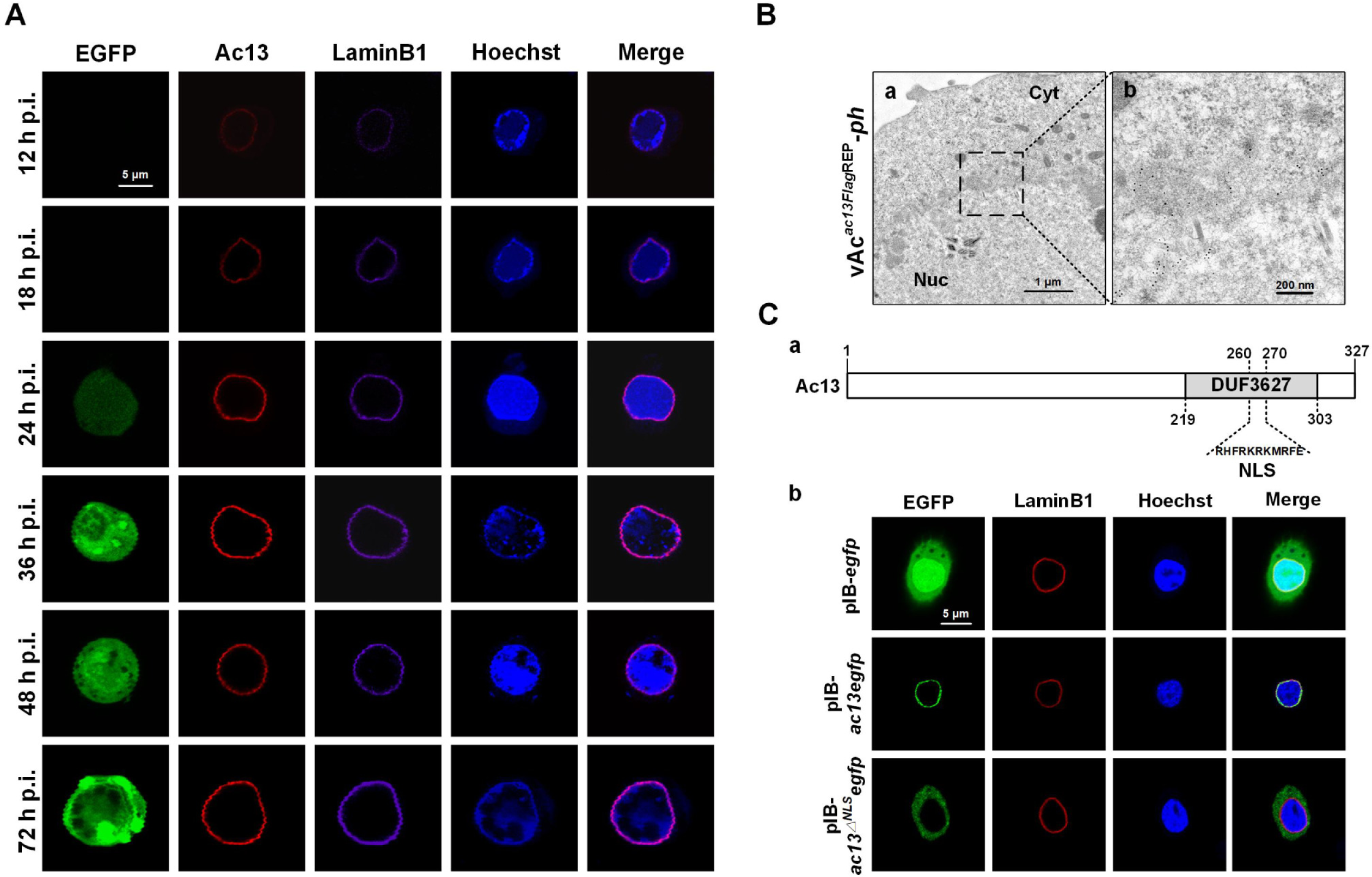
Subcellular localization of Ac13 in Sf9 cells. (A) Immunofluorescence analysis. Sf9 cells, infected with vAc^*ac13Flag*REP^-*ph* at an MOI of 5, were fixed with paraformaldehyde at the indicated time points and immunostained with an anti-FLAG antibody to detect Ac13 (red), an anti-Dm0 antibody to detect LaminB1 (purple). EGFP was an indicator of cells infected with virus (green). The nuclei were stained with Hoechst33258 (blue). Bars, 5 μm. (B) Immunoelectron microscopy analysis. Sf9 cells were infected with vAc^*ac13Flag*REP^-*ph* at an MOI of 10 and harvested at 48 h p.i. The ultrathin sections were probed with anti-FLAG antibody as the primary antibody and goat anti-rabbit IgG coated with gold particles (10 nm) as the secondary. Bars, 1 μm and 200 nm. (C) (a) Schematic representation of the NLS of Ac13. The NLS of Ac13 was predicted in the DUF3627. (b) Fluorescence microscope analysis. Sf9 cell, transfected with the plasmids pIB-*ac13egfp*, 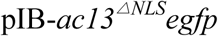 or pIB-*egfp*, were analyzed by immunofluorescence microscopy at 24 h p.t.. EGFP was an indicator of Ac13 (green), and an anti-Dm0 antibody was used to detect the LaminB1 in cells (red). The nuclei were stained with Hoechst 33258 (blue). Bars, 5 μm.

It was previously reported that Bm5 localized to the nuclear membrane in a DUF3627-dependent manner (32). Bioinformatic analysis indicated that there was a putative nuclear localization signal (NLS) motif in the Ac13 DUF3627 domain (Fig. 2C a). To further examine the location of Ac13 in the absence of viral infection or the NLS motif, Sf9 cells were transfected with the pIB-*ac13egfp*, 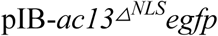 or pIB-*egfp* plasmids. As shown in Fig. 2C b, pIB-*ac13egfp* fluorescence was detected at the nuclear membrane, whereas 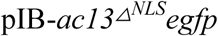, encoding a disrupted NLS, fluorescence was observed only in the cytoplasm. As a control, pIB-*egfp* showed homogenous fluorescence throughout the cytoplasm and nucleus. These results indicated that Ac13 located to the nuclear membrane independently of viral infection, but nuclear import required the NLS motif of DUF3627.

### 3. Ac13 is essential for BV production but not for OB formation

To investigate the function of *ac13* in the AcMNPV life cycle, an *ac13*-null bacmid, bAc^*ac13*KO^-*ph*, an *ac13*-rescue bacmid, bAc^*ac13*REP^-*ph*, and a pseudo-wild-type bacmid, bAc-*ph* (Fig. 3), were constructed. All bacmids were confirmed by PCR (primer pairs shown in Table 1) and DNA sequencing (data not shown).

**Table 1.**
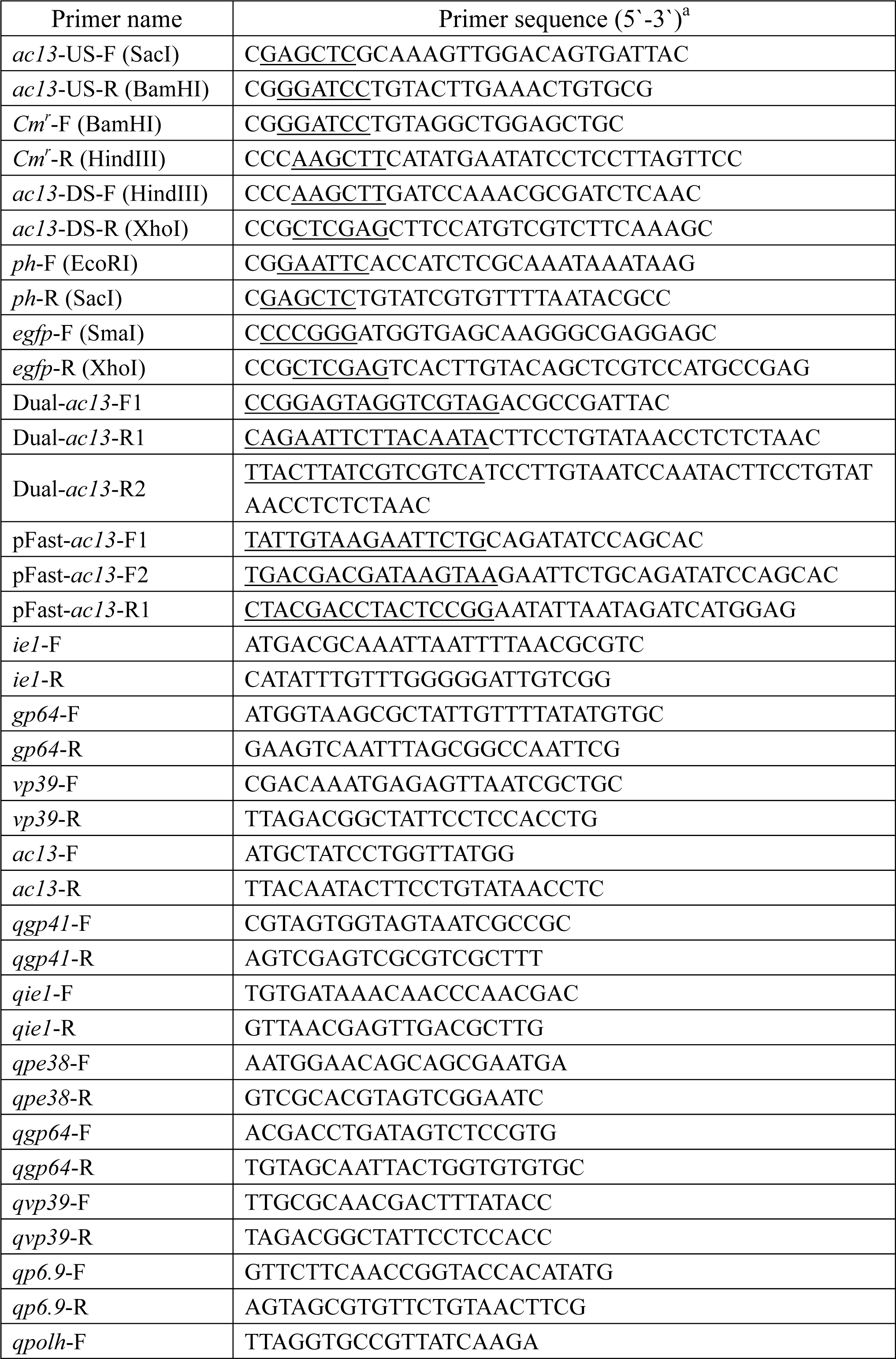
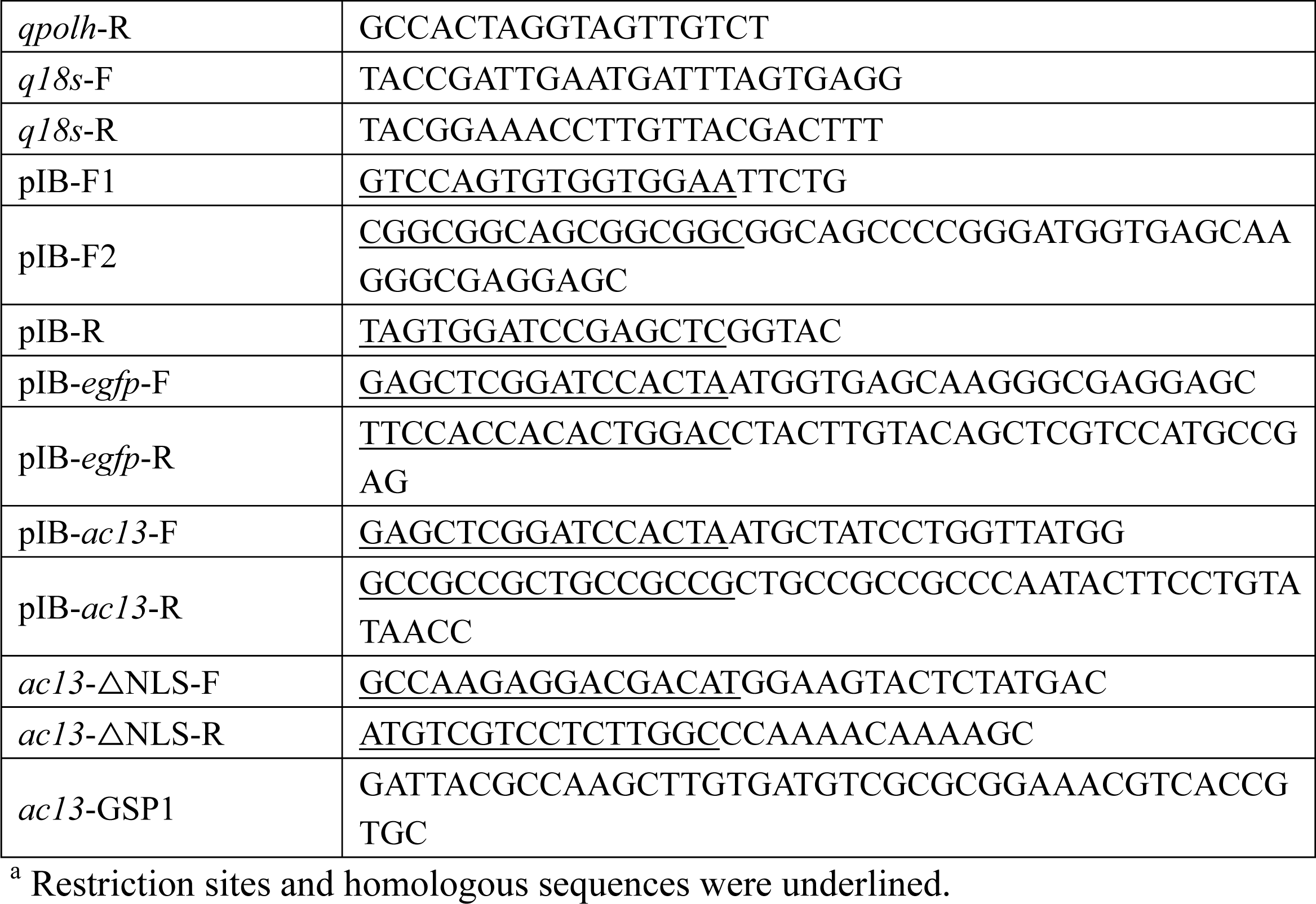
Primers used in this study.

**FIG 3.**
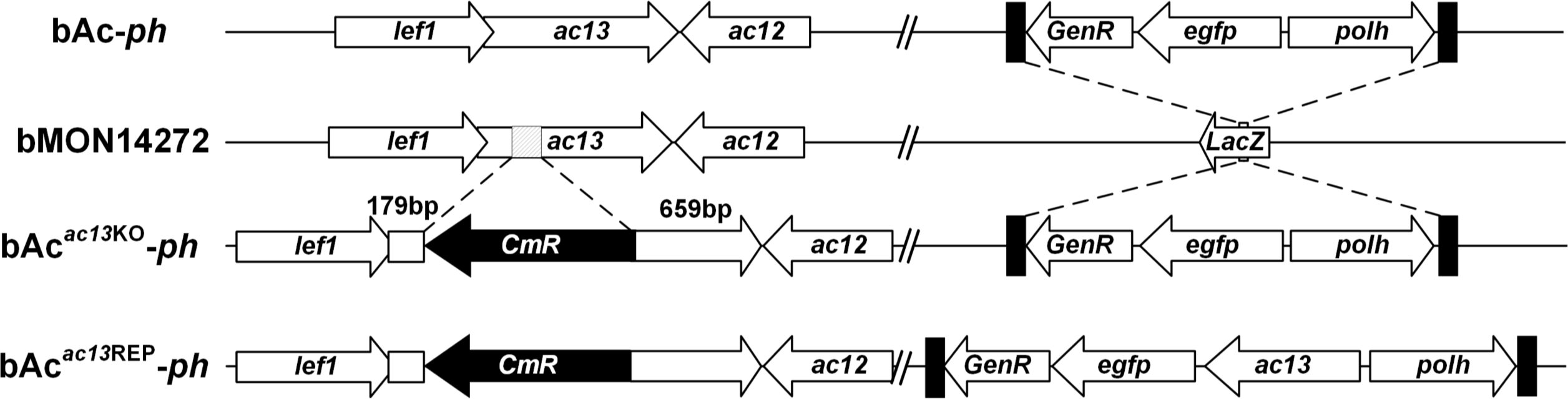
Schematic diagram of bAc^*ac13*KO^-*ph*, bAc^*ac13*REP^-*ph* and bAc-*ph* construction. Using the bacmid bMON14272, the bAc13KO was generated by replacing 146 bp fragment of the *ac13* ORF with a chloramphenicol resistance (*Cm*^r^) gene cassette via homologous recombination. The *egfp* and *polh* genes were inserted into the *polh* locus of bAc13KO via Tn7-mediated transposition to generate bAc^*ac13*KO^-*ph*. The *ac13* together with the *egfp* and *polh* genes were inserted into the *polh* locus of bAc13KO to generate bAc^*ac13*REP^-*ph*. bAc-*ph* was constructed by inserting the *egfp* and *polh* genes into the *polh* locus of bMON14272.

To determine the effects of the absence of *ac13* on viral proliferation and OB morphogenesis, Sf9 cells were transfected with the bAc^*ac13*KO^-*ph*, bAc^*ac13*REP^-*ph* or bAc-*ph* bacmids and monitored under a fluorescent microscope. No obvious differences were found in the numbers of fluorescent cells between the three bacmids at 24 h p.t. (Fig. 4A, upper row), indicating that the three bacmids had comparable transfection efficiencies. By 96 h p.t., almost all cells transfected with bAc^*ac13*REP^-*ph* or bAc-*ph* showed fluorescence, whereas the number of fluorescent bAc^*ac13*KO^-*ph*-transfected cells increased only slightly from 24 to 96 h p.t. (Fig. 4A, upper and middle rows). This result indicated that although bAc^*ac13*KO^-*ph*-transfected cells were able to produce infectious BVs, the efficiency of BV production was impaired.

**FIG 4.**
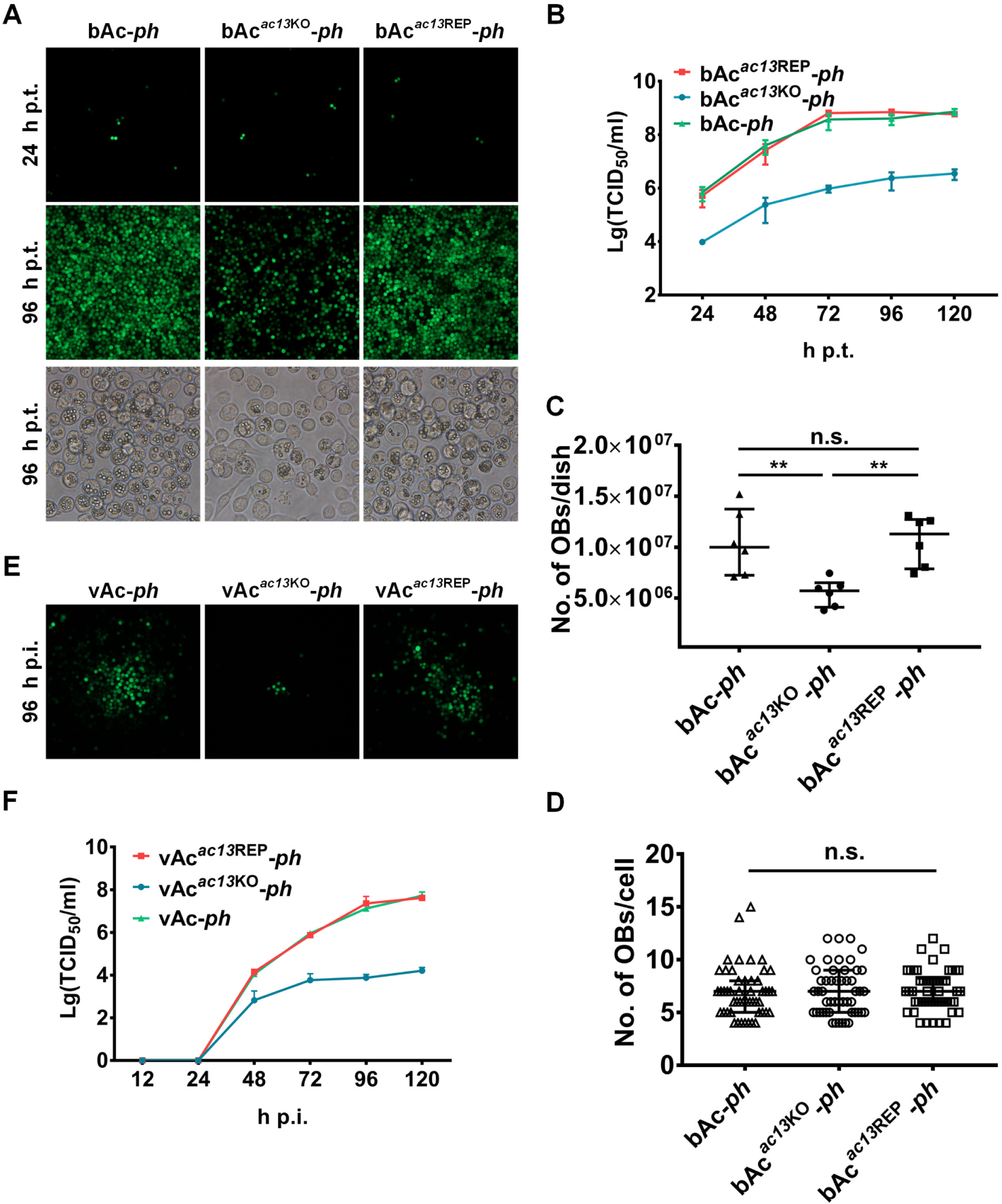
Analysis of viral replication and occlusion body formation in the transfected/infected Sf9 cells. (A) Fluorescence microscopy of cells transfected with the bacmids of bAc-*ph*, bAc^*ac13*KO^-*ph* or bAc^*ac13*REP^-*ph* at 24 or 96 h p.t. (upper and middle rows). Light microscopy of cells transfected with each bacmid at 96 h p.t. (lower row). (B) The supernatants of Sf9 cells, transfected with bAc-*ph*, bAc^*ac13*KO^-*ph* or bAc^*ac13*REP^-*ph*, were harvested at the designated time points and quantified by TCID_50_ endpoint dilution assays. Each data points represent average titers from three separate transfections. Error bars represent standard deviations (SD). (C) OB production in each dish. The cells were gently scraped, and total OBs were measured using hemocytometer at 96 h p.t.. ** indicates *p* < 0.01, n.s. indicates no significance, *p* > 0.05. (D) OB production in each cell. The number of envisaged-visible OBs was under phase contrast microscope counted at 96 h p.t., and more than 50 cells were counted for each condition. n.s. indicates no significance, *p* > 0.05. (E) Fluorescence microscopy images of Sf9 cells infected with vAc-*ph*, vAc^*ac13*KO^-*ph* or vAc^*ac13*REP^-*ph* at an MOI of 0.002 at 96 h p.i.. (F) Sf9 cells were infected with vAc-*ph*, vAc^*ac13*KO^-*ph* or vAc^*ac13*REP^-*ph* at an MOI of 0.002. The supernatants were collected at the indicated time points and determined by TCID_50_ endpoint dilution assays. Each data points represent average titers from three separate infections. Error bars represent SD.

To further examine the effects of *ac13* deletion on viral proliferation, we collected BV supernatants from cells transfected with each bacmid at the indicated time points, determined BV titers using the 50% tissue culture infective dose (TCID_50_) endpoint dilution assay, and performed a viral growth curve analysis. Viruses produced from the bAc^*ac13*REP^-*ph* and bAc-*ph* bacmids showed similar growth kinetics, reaching 7.0×10^8^ and 4.0×10^8^ TCID_50_/mL at 96 h p.t., respectively. However, the TCID_50_ of bAc^*ac13*KO^-*ph* was reduced by 400-fold compared with bAc^*ac13*REP^-*ph* and bAc-*ph* at 120 h p.t. (*p* < 0.001) (Fig. 4B). Light microscopy analysis revealed that a large proportion of cells transfected with bAc-*ph* and bAc^*ac13*REP^-*ph* contained OBs at 96 h p.t., whereas only a small proportion of bAc^*ac13*KO^-*ph*-transfected cells contained OBs (Fig. 4A, lower panel). Subsequently, we collected cells transfected with bAc^*ac13*KO^-*ph*, bAc^*ac13*REP^-*ph* or bAc-*ph* from each dish at 96 h p.t. and counted OBs using a hemocytometer. As shown in Fig. 4C, the numbers of OBs produced by cells transfected with bAc^*ac13*KO^-*ph* were significantly reduced compared with the numbers produced by cells transfected with bAc-*ph* and bAc^*ac13*REP^-*ph*. These results indicated that BV production in bAc^*ac13*KO^-*ph*-transfected cells was significantly reduced compared with cells transfected with bAc-*ph* and bAc^*ac13*REP^-*ph*. By contrast, no obvious differences were found in the average numbers of OBs produced by each cell transfected with bAc^*ac13*KO^-*ph*, bAc^*ac13*REP^-*ph* or bAc-*ph* at 96 h p.t. (Fig. 4A, lower panel and Fig. 4D), indicating that *ac13* deletion had no effect on OB formation.

To confirm the results obtained following bacmid transfection, Sf9 cells were infected with vAc^*ac13*KO^-*ph*, vAc^*ac13*REP^-*ph* or vAc-*ph* at an MOI of 0.002 and a viral growth curve analysis was performed. As shown in Fig. 4E and F, vAc^*ac13*REP^-*ph* showed similar growth kinetics to vAc-*ph*. However, BV production in vAc^*ac13*KO^-*ph*-infected cells was significantly reduced compared with BV production in cells infected with vAc-*ph* and vAc^*ac13*REP^-*ph*. Taken together, these data suggested that *ac13* deletion significantly decreased BV production, but did not affect OB formation.

### 4. *ac13* is not required for synthesis of viral DNA or transcription of viral genes

The BV life cycle includes replication of viral DNA, assembly of progeny nucleocapsids, egress from the nucleus, transport through the cytoplasm, and budding from the plasma membrane where the BV gains its envelope. To determine whether reduced vAc^*ac13*KO^-*ph* BV production resulted from a defect in viral DNA replication, viral DNA replication was compared between bAc^*ac13*KO^-*ph-* and bAc-*ph*-transfected cells by quantitative PCR (qPCR) within 24 h p.t., before secondary infection by BVs could occur (15). Sf9 cells were transfected with equal amounts of bAc^*ac13*KO^-*ph* or bAc-*ph* bacmid DNA and collected at 0, 12 and 24 h p.t.. Total intracellular DNA was extracted and treated with DpnI to eliminate input bacmid DNA. The viral genome copy number was measured by qPCR using *gp41*-specific primers as previously reported (33). As shown in Fig. 5A, the viral DNA content in bAc^*ac13*KO^-*ph* and bAc-*ph-*transfected cells both increased with similar rates from 0 to 24 h p.t. (*p* > 0.05), indicating that *ac13* deletion did not affect viral DNA replication. Subsequently, expression of six viral genes (*ie1, pe38, gp64, vp39, p6*.*9* and *polh*) was analyzed by reverse transcription (RT)-qPCR. As shown in Fig. 5B, no significant differences were found in the transcript levels of any genes between bAc^*ac13*KO^-*ph*-and bAc-*ph*-transfected cells (*p* > 0.05). These results suggested that *ac13* deletion did not affect early or late viral gene transcription.

**FIG 5.**
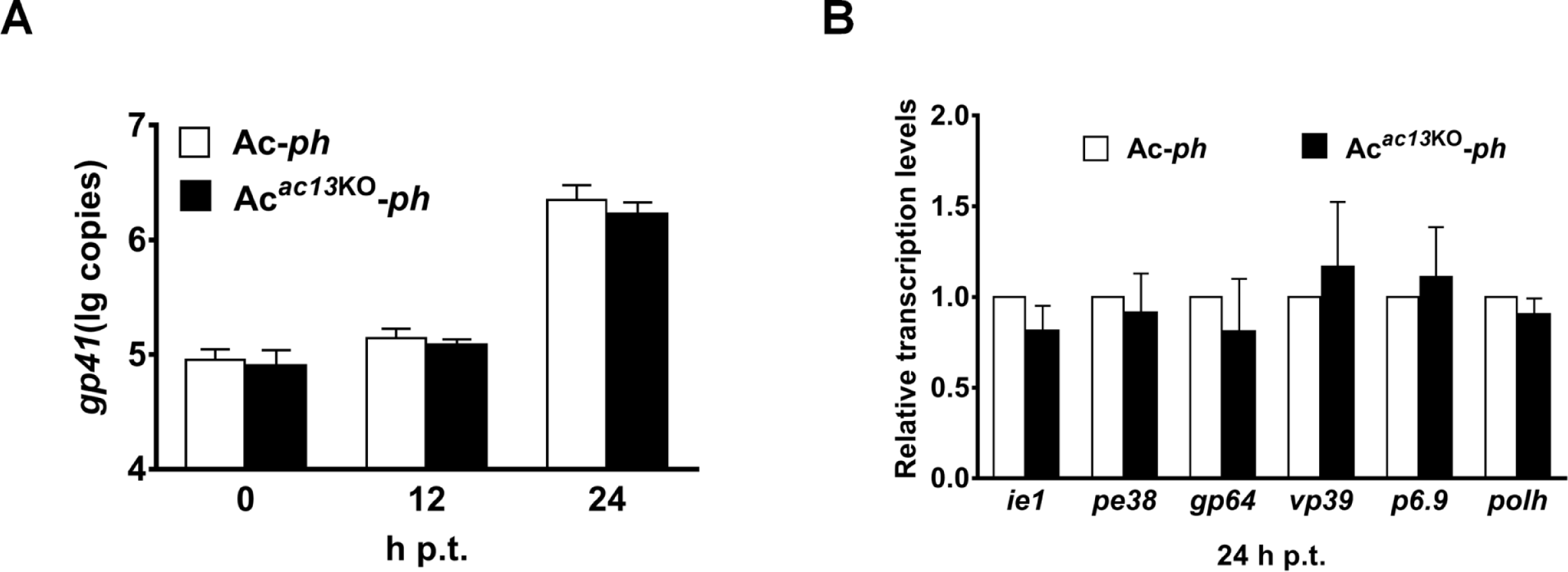
Analysis of viral DNA replication and viral genes transcription. (A) qPCR analysis of viral DNA replication. Total cellular DNAs were extracted at the indicated time points from Sf9 cells transfected with the bacmids of bAc^*ac13*KO^-*ph* or bAc-*ph* and were digested with restriction enzyme DpnI to eliminate input bacmid DNA. The genomes copies were analyzed with qPCR using the *gp41*-specific primer pairs. The values represent the averages from three independent assays. Error bars indicate SD. (B) RT-qPCR analysis of viral genes transcription at 24 h p.t.. Total cellular RNA was extracted at 24 h p.t. from Sf9 cells which were transfected with bAc^*ac13*KO^-*ph* or bAc-*ph*. The transcription of selected viral genes was measured with RT-qPCR. The transcript level of each gene was normalized to that of the cell 18S rRNA. The values represent the averages from three independent assays, and error bars represent SD.

### 5. Ac13 is required for efficient nuclear egress

To further explore the impediments to BV production in the absence of *ac13*, transmission electron microscopy (TEM) was used to examine thin sections generated from cells infected with vAc^*ac13*KO^-*ph*, vAc^*ac13*REP^-*ph* or vAc-*ph* at an MOI of 10 at 48 h p.i.. As shown in Fig. 6A, the typical symptoms of baculovirus infection appeared both in vAc^*ac13*KO^-*ph*-and vAc-*ph*-infected cells. These included an enlarged nucleus with a net-shaped VS, a large number of rod-shaped electron-dense nucleocapsids within the VS, and mature ODVs with multiple nucleocapsids and ODV-containing OBs around the ring zone. As expected, vAc^*ac13*REP^-*ph*-infected cells exhibited similar characteristics to those of cells infected with vAc-*ph*. According to the above results, *ac13* deletion did not affect either nucleocapsid assembly or OB formation.

**FIG 6.**
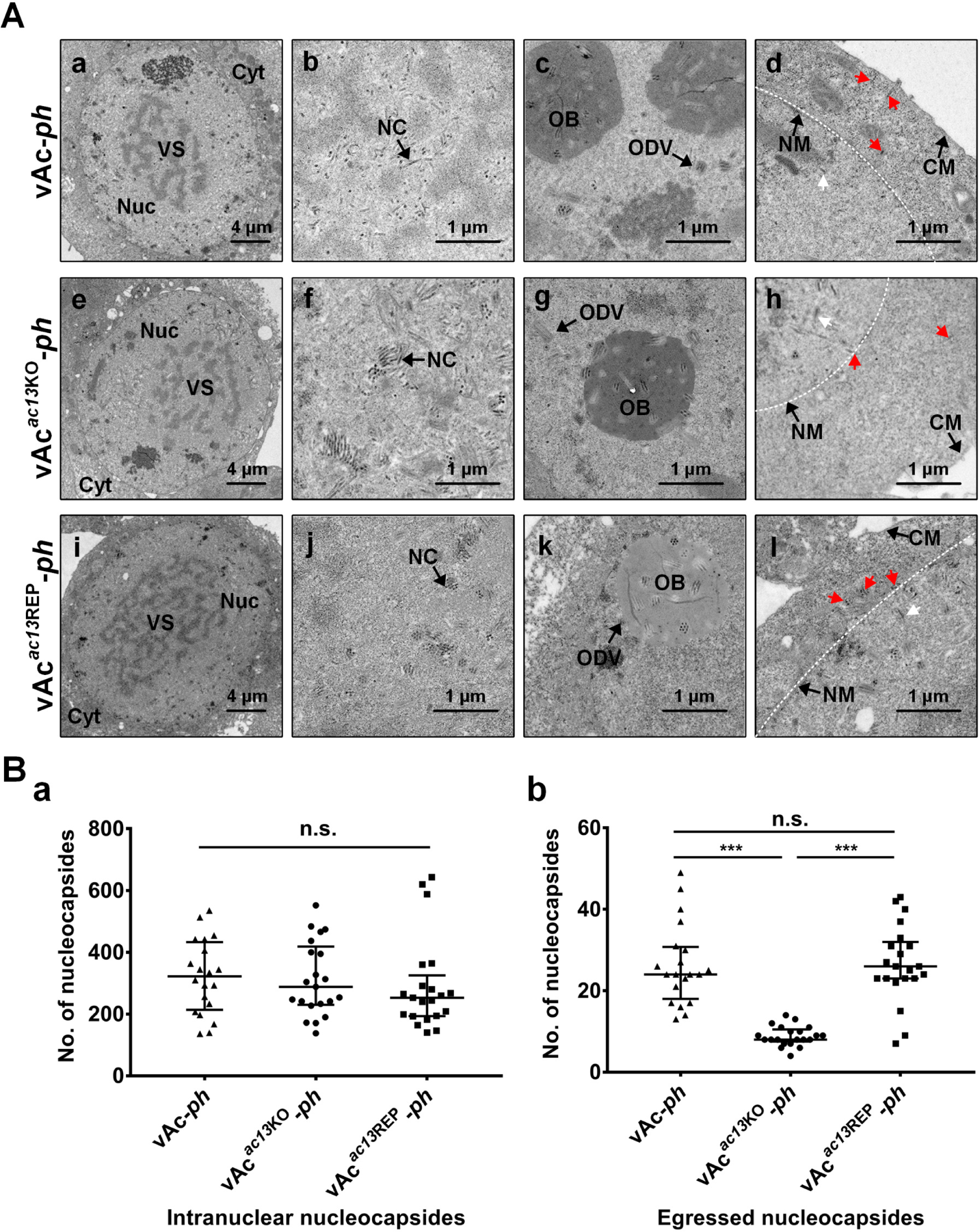
Transmission electron microscopy analyses of Sf9 cells infected with vAc-*ph*, vAc^*ac13*KO^-*ph* and vAc^*ac13*REP^-*ph*. (A) Sf9 cells, infected with vAc-*ph*, vAc^*ac13*KO^-*ph* or vAc^*ac13*REP^-*ph* at an MOI of 10, were fixed at 48 h p.i. and prepared for TEM. (a to d) Cells infected with vAc-*ph*. (e to h) Cells infected with vAc^*ac13*KO^-*ph*. (i to l) Cells infected with vAc^*ac13*REP^-*ph*. (a, e and i) Enlarged nucleus (Nuc) and virogenic stroma (VS) in cells infected with vAc-*ph*, vAc^*ac13*KO^-*ph* or vAc^*ac13*REP^-*ph*. (b, f and j) Normal electron-dense nucleocapsids (NC) in cells infected with vAc-*ph*, vAc^*ac13*KO^-*ph* or vAc^*ac13*REP^-*ph*. (c, g and k) ODVs containing multiple nucleocapsids and OBs embeding normal ODVs in cells infected with vAc-*ph*, vAc^*ac13*KO^-*ph* or vAc^*ac13*REP^-*ph*. (d, h and l) Nucleocapsids residing in the cytoplasm or budding from the nuclear or cytoplasmic membranes were indicated with red arrows, while nucleocapsids residing in the nucleus were indicated with white arrows in cells. The nuclear membrane was shown with white dotted line. Bars, 2 μm (a,e and i) and 500 nm (b to d, f to h and j to l). Nuc, nucleus; Cyt, cytoplasm; NM, nuclear membrane; CM, cytoplasmic membrane. (B) The numbers of intranuclear nucleocapsids (n.s. indicates no significance, *p* > 0.05) (a) and egressed nucleocapsids (*** indicates *p* < 0.001, n.s. indicates no significance, *p* > 0.05) (b) were determined. Numbers were calculated from 20 cells.

Ac13 localized to the nuclear membrane, and therefore it might play a role in egress of nucleocapsids from the nucleus. The TEM analysis was used to assess whether *ac13* deletion had any effect on egress. According to methods previously reported (15, 16, 24), we counted and compared the numbers of intranuclear and egressed nucleocapsids in 20 randomly chosen cells infected with vAc^*ac13*KO^-*ph*, vAc^*ac13*REP^-*ph* or vAc-*ph*. The egressed nucleocapsids included nucleocapsids exiting the nuclear membrane, in transport through the cytoplasm and budding from the cytoplasmic membrane (Fig. 6A d, h and i). Intranuclear nucleocapsids of vAc^*ac13*KO^-*ph*-infected cells were comparable to those of vAc^*ac13*REP^-*ph*-and vAc-*ph*-infected cells (Fig. 6B a). By contrast, egressed nucleocapsids were substantially reduced in vAc^*ac13*KO^-*ph*-infected cells compared with vAc^*ac13*REP^-*ph*-and vAc-*ph*-infected cells (*p* < 0.001) (Fig. 6B b). Taken together, these data showed that *ac13* deletion impaired nucleocapsid egress.

### 6. *ac13* deletion does not affect OB morphogenesis in *Spodoptera exigua* larvae

The above results indicated that *ac13* deletion did not affect the number of OBs formed within each cell. To further investigate whether the absence of *ac13* had an effect on OB morphogenesis in larvae, scanning electron microscopy (SEM) and TEM were performed on OBs purified from *S. exigua* cadavers infected with vAc-*ph*, vAc^*ac13*KO^-*ph* or vAc^*ac13*REP^-*ph*. As shown in Fig. 7A, the OBs formed within vAc^*ac13*KO^-*ph*-infected larvae had smooth surfaces and sharp edges and contained normal ODVs, similar to those of vAc-*ph*-or vAc^*ac13*REP^-*ph*-infected larvae. Subsequently, we negatively stained and counted the numbers of ODVs within OBs of the three viruses using TEM. The average number of ODVs per OB of Ac^*ac13*KO^-*ph* was comparable to those of vAc-*ph* or vAc^*ac13*REP^-*ph* (*p* > 0.05) (Fig. 7B). Taken together, these results showed that *ac13* was not required for OB morphogenesis in larvae.

**FIG 7.**
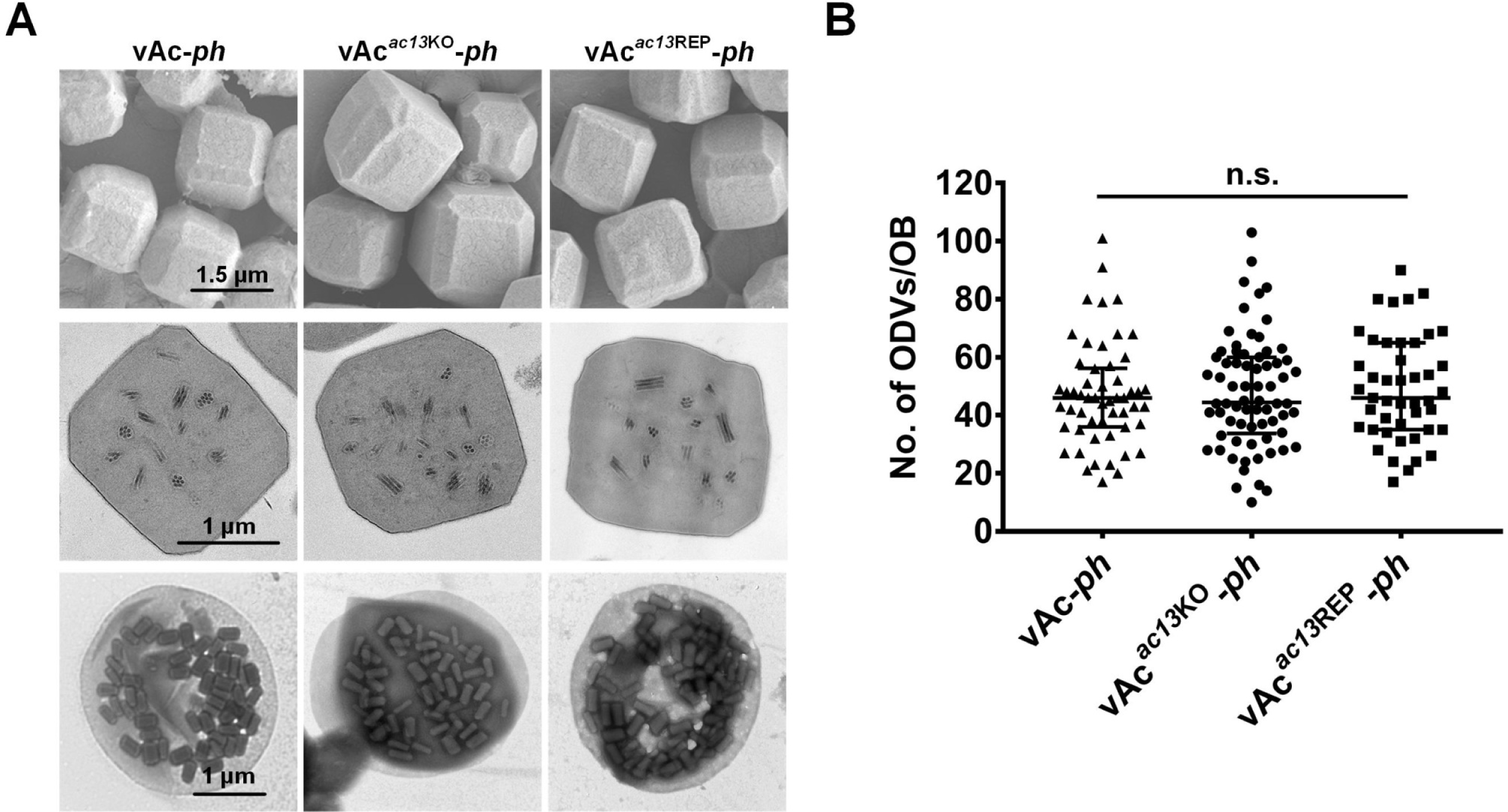
Scanning electron microscopy, transmission electron microscopy and negative staining analysis of OBs from vAc-*ph*, vAc^*ac13*KO^-*ph* and vAc^*ac13*REP^-*ph*. (A) The OBs, purified from *S. exigua* larvae infected with vAc-*ph*, vAc^*ac13*KO^-*ph* or vAc^*ac13*REP^-*ph*, were observed with Scanning electron microscopy (upper row), transmission electron microscopy (middle row) and negative staining after treating by dissolution buffer on the grid (lower row). (B) Numbers of ODVs embedded in each OB. More than 40 OBs of each virus were analyzed. n.s. indicates no significance (*p* > 0.05).

## DISCUSSION

The *ac13* gene is conserved in all sequenced alphabaculoviruses, implying that it may play an important role in the viral life cycle. In the present study, we investigated the role of *ac13* in AcMNPV by constructing an *ac13*-null bacmid (bAc^*ac13*KO^-*ph*). We determined that *ac13* was required for efficient egress of nucleocapsids from the nucleus to the cytoplasm, but not for OB formation.

The 5’ RACE and sequence analyses indicated that *ac13* was regulated by an atypical early promoter (GCAGT) and a canonical late promoter (TAAG) in AcMNPV-infected Sf9 cells. The TSS of the late promoter was consistent with previous studies. However, the TSS of the early promoter differed by 2 nt from previous data generated from AcMNPV-infected *Trichoplusia ni* cells. A potential explanation for this discrepancy might be that different hosts impact the TSS usage of AcMNPV. In addition to *ac13*, 11 other genes (*ac23, pkip, v-fgf, pp31, odv-e66, ac82, he65, gp64, p35, me53* and *ie0*) contain both early (TATAA or CAGT) and late (TAAG) motifs in AcMNPV (27). Global analysis of AcMNPV gene expression in the midgut of *T. ni* indicated that *ac13* gene transcripts could be detected from 6 to 72 h p.i, with higher expression than most other AcMNPV genes (34). Our analysis of *ac13* temporal transcription patterns agreed with those results. Thus, *ac13* may play an important role in infection *in vivo* and *in vitro* both at early and late stages, although the Ac13 protein can only be detected during late infection. However, we cannot rule out the possibility that levels of Ac13 expression during the early stage were too low to be detected. It is also possible that the *ac13* gene may play a role in early infection not through an encoded protein but through other types of gene products (i.e., peptides or non-coding RNAs), or may participate as a DNA element. Several *cis*-acting elements have been identified in AcMNPV. For example, Ac83 was required for *per os* infectivity factor (PIF) complex formation and was deemed a true PIF (35), while *ac83* is involved in nucleocapsid envelopment via an internal *cis*-acting element (36). We found that Ac13 localized at the nuclear membrane independently of viral infection, consistent with a prior study of the subcellular localization of Bm5 (32). Protein nuclear import typically requires a NLS, which assists transport through the nuclear pore complex into the nucleus. Previous studies identified several proteins with NLSs in AcMNPV including LEF-3, DNApol, IE1 and BV/ODV-C42 (37-39). Bioinformatic and confocal microscopy analyses indicated the presence of an NLS motif in Ac13, which played an essential role in its nuclear import (Fig. 2C). However, bioinformatic analysis suggested that Ac13 is not an integral membrane protein. Yet-unknown proteins may facilitate the nuclear membrane accumulation of Ac13.

It was reported that *bm5* encodes a multifunctional protein that regulates viral transcription and OB formation and contributed to BV production (32). However, in our study, observation of cells transfected with bAc^*ac13*KO^-*ph* and bAc-*ph* demonstrated that *ac13* deletion reduced BV production by 99.7%, but did not affect OB formation. The *Cm*^*r*^ cassette was reversibly inserted in the *ac13* ORF, completely disrupting *ac13* expression without affecting transcription or expression of the neighbor genes *lef1* and *ac12*. The phenotype of the *ac13* knockout could be rescued by a repair virus. Similarly, a repair virus was able to rescue the phenotype associated with *bm5* deletion (35). Thus, OB formation was affected by *bm5* deletion, whereas OBs were observed in cells transfected with bAc^*ac13*KO^-*ph* at near-wild type levels (Fig. 4 and Fig. 7). This discrepancy may result from the different measurement methods use. In the Bm5 study, OB formation was confirmed by counting the number of OBs in the whole dish. Because *bm5* deletion reduced BV production and decreased the number of cells undergoing secondary infection, the number of *bm5*-null*-*infected cells may have been lower than the number of cells infected with wild-type virus. In our study, we assessed OB formation by counting the number of OBs per cell and observed the shape and inner structure of OBs via SEM and TEM. Differences may also have resulted from the different viruses used. Several genes have been reported to show discrepancies in knockout phenotypes between AcMNPV and BmNPV. For instance, *gp41* is essential for AcMNPV replication (21) but not for BmNPV replication (30). Additionally, *ac51* is required for BV production but not for OB morphogenesis in AcMNPV (15), while *bm40*, an orthologous gene of *ac51* in BmNPV, is essential for BV and OB morphogenesis (40).

The nuclear membrane accumulation of Ac13 suggested that it may be involved in nucleocapsid egress from the nucleus. In the alphaherpesviruses, pUL31 and pUL34, which are required for egress of nucleocapsids from the nucleus to cytoplasm, are codependently localized to the nuclear rim (41). Largely based on the results of TEM analyses, nucleocapsid egress from the nucleus is thought to occur through a process of budding from the nuclear membrane (42), and a recent study demonstrated that baculovirus nucleocapsid egress from the nucleus by disrupting the nuclear membrane (12). We used TEM to examine and compare cells infected with vAc^*ac13*KO^-*ph*, vAc^*ac13*REP^-*ph* and vAc-*ph*. The results showed that *ac13* deletion did not affect nucleocapsid assembly or progression of viral infection into very late phases to form OBs, but impaired efficient egress of nucleocapsids from the nucleus to the cytoplasm. Several AcMNPV genes have been reported to be involved in nuclear egress of nucleocapsids including *ac11, ac51, ac66, ac75, ac78, gp41, ac93, p48, exon0* and *ac142*. The first to be identified, *exon0*, is not strictly essential for production of BVs because a few nucleocapsids in cells infected with an *exon0*-null virus did pass through the nuclear membrane. Recently, Qiu et al. demonstrated that the *ac51* deletion affected the efficiency of nuclear egress of nucleocapsids, but did not affect nucleocapsid assembly and ODV envelopment (15). In this study, the genes required for the nuclear egress of nucleocapsids and production of BVs seemed to be divided into two categories (15): (*i*) genes whose deletion did not affect nucleocapsid assembly but prevented nuclear egress of nucleocapsids, thus abrogating BV production and also interrupting OB formation, and (*ii*) genes whose deletion did not affect nucleocapsid assembly but decreased the efficiency of nuclear egress of nucleocapsids, thus decreasing BV production but not affecting OB formation. According to previous reports, *exon0,ac66* and *ac51* belong to the second category. Our results confirm that *ac13* is a fourth gene belonging to the second category.

In summary, nucleocapsid egress is essential for mature BV production and virus proliferation. Although many host and viral genes associated with nuclear egress have been determined, the process is incompletely understood in baculoviruses. In the present study, we investigated the functions of *ac13* and found that it was required for efficient nuclear egress of nucleocapsids during BV production, but not for OB formation *in vivo* and *in vitro*.

## MATERIALS AND METHODS

### Cells, viruses, insect and antibodies

Sf9 cells were cultured at 27°C in Grace’s medium (Invitrogen, Carlsbad, CA, USA) supplemented with 10% fetal bovine serum (Gibco, Grand Island, NY, USA). AcMNPV recombinant bacmids were constructed with the bMON14272 bacmid (Invitrogen) and maintained in *Escherichia coli* strain DH10B (Invitrogen), which also contained helper plasmids for homologous recombination and transposition.

The anti-Ac13 polyclonal antiserum was prepared in rabbits according to previously published methods (21). The polyclonal anti-GP64 and anti-VP39 antibodies were gifts from Z. H. Hu (Wuhan Institute of Virology, China). Mouse monoclonal anti-actin antibody was from Proteintech (Wuhan, China) and mouse monoclonal anti-Dm0 antibody was from the Developmental Studies Hybridoma Bank (DSHB; Iowa City, IA, USA).

### Time course analysis of transcription and expression

Sf9 cells (1.0×10^6^ cells/F35-mm plate) were infected with AcMNPV at an MOI of 5 and collected at 0, 3, 6, 12, 24, 36 and 48 h p.i.. Total RNA was extracted using RNAiso Plus (TaKaRa, Dalian, China) according to the manufacturer’s instructions and quantitated using a Nanodrop-2000 spectrophotometer (Thermo Fisher Scientific, Waltham, MA, USA). Subsequently, cDNA was synthesized using an iScript cDNA synthesis kit (Bio-Rad, Hercules, CA, USA). Finally, transcripts were detected by PCR using gene-specific primer pairs (primer sequences shown in Table 1) and the above cDNA as a template.

For the time course analysis of Ac13 expression, Sf9 cells (1.0×10^6^ cells/F35-mm plate) were infected with AcMNPV at an MOI of 10 and harvested at 0, 6, 12, 18, 24 and 48 h p.i.. Subsequently, western blotting was performed with anti-Ac13 (1:1,000) primary antibody and horseradish peroxidase-conjugated goat anti-rabbit (1:3,000; Proteintech) secondary antibody. A protein ladder (Thermo Fisher Scientific) was used to judge protein sizes.

### The 5’ RACE analysis

Sf9 cells (1.0×10^6^ cells/F35-mm plate) were infected with AcMNPV at an MOI of 5 and collected at 4 and 24 h p.i. Total RNA was isolated using RNAiso Plus (TaKaRa). The 5’ RACE reaction was performed with an *ac13*-specific primer (*ac13*-GSP1) using a SMARTer^®^ RACE 5’/3’ Kit (TaKaRa) according to the manufacturer’s protocol. The PCR products were cloned into the pMD19-T vector (TaKaRa) and sequenced.

### Construction of *ac13* knockout and repaired bacmids

An *ac13* knockout bacmid was constructed as previously described (43, 44). First, a 618-bp sequence upstream and a 686-bp sequence downstream of the *ac13* ORF were amplified by PCR using the primer pairs *ac13-*US-F/R and *ac13-*DS-F/R, respectively, and the AcMNPV bacmid as template. A 1,137-bp fragment was amplified using the primer pairs *Cm*^*r*^-F/R using the pUC18-*Cm*^*r*^ plasmid as the template. Subsequently, the 618-bp upstream fragment, the 686-bp downstream fragment and the 1,137-bp fragment were double-digested with SacI/BamHI, HindIII/XhoI and BamHI/HindIII, respectively. The three restriction digestion fragments were gel purified and consecutively ligated into the pBlueScript II SK(+) vector to generate pSK-*ac13*US-*Cm*^*r*^-*ac13*DS. A fragment, amplified using the primer pair *ac13-*US-F/*ac13-*DS-R and template pSK-*ac13*US-*Cm*^*r*^-*ac13*DS, was used to electroporate *E. coli* BW25113 cells (containing bMON14727 and pKD46) to replace the N-terminal 174-bp (nt 43514 nt to 43638 of the AcMNPV genome) of *ac13* with the *Cm*^*r*^ cassette via λRed homologous recombination. The resulting *ac13-*null bacmid, confirmed by PCR and DNA sequencing, was named bAc^*ac13*KO^.

Subsequently, the *polh* and the *egfp* genes were separately cloned into the pFastBacDual vector (Invitrogen) under the control of the *polh* and *p10* promoters via restriction digestion and ligation to generate pFBD-*ph*-*egfp*. DH10B competent cells, containing the helper plasmid pMON7124 and the bAc^*ac13*KO^ bacmid, were transfected with the pFBD-*ph*-*egfp* plasmid, generating an *ac13*-null bacmid (bAc^*ac13*KO^-*ph*) by Tn7-mediated transposition. Similarly, a wild-type control bacmid (bAc-*ph*) was generated by inserting the *polh* and *egfp* genes into the *polh* locus of bMON1427. To construct an *ac13* rescue bacmid, a 1,458-bp fragment containing the *ac13* native promoter and ORF was amplified by PCR with the primer pair Dual-*ac13*-F1/R1 from the bMON1427 template. The 1,458-bp fragment was inserted in the pFBD-*ph*-*egfp* plasmid to produce pFBD-*ph*-*ac13*-*egfp* via homologous recombination. This vector was then used to transform DH10B competent cells (containing bAc^*ac13*KO^ and pMON7124) to generate an *ac13*-rescue bacmid (bAc^*ac13*REP^-*ph*). Meanwhile, another *ac13* rescue bacmid bAc^*ac13Flag*REP^-*ph*, encoding a FLAG tag at its 3’-end, was constructed using the same method. All recombinant bacmids were confirmed by PCR and DNA sequencing.

### Construction of *ac13* subcellular localization plasmids

The transient expression plasmid pIB-*egfp* was constructed using FastCloning (45). Briefly, the pIB/V5-His vector (Invitrogen) and insert *egfp* fragment were amplified by PCR. The *egfp* fragment 16 bp sequence was homologous with the vector. The PCR products were digested with DpnI (TaKaRa) at 37°C for 1 h, and then used to transform *E. coli* DH5*α* competent cells. Subsequently, the *ac13* ORF was amplified from the AcMNPV bacmid and subcloned into pIB-*egfp* in-frame with the *egfp* fragment to generate pIB-*ac13egfp* by FastCloning. Based on the pIB-*ac13egfp* vector, the 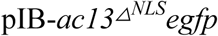 vector bearing a truncated *ac13* gene with an NLS deletion (aa 778–810) was also constructed by FastCloning.

### Transfection and infection assay

Sf9 cells were seeded in a 35-mm diameter six-well plate (1.0×10^6^ cells/well) and allowed to attach for 2 h at 27°C. The cells were transfected in triplicate with 10 μg of each bacmid DNA (bAc^*ac13*KO^-*ph*, bAc^*ac13*REP^-*ph* and bAc-*ph*)using 8 μL of Cellfectin II (Invitrogen) according to the manufacturer’s instructions. The transfection buffer was then replaced with fresh Grace’s medium after incubation for 5 h. The BVs contained in the supernatant was called vAc^*ac13*KO^-*ph*, vAc^*ac13*REP^-*ph*, and vAc-*ph*. BVs were harvested at 24, 48, 72, 96 and 120 h p.t. and viral titers were determined using a TCID_50_ endpoint dilution assay. Cells were infected in triplicate with each virus at an MOI of 0.002. After viral absorption for 1 h at 27°C, the infection mixture was replaced with fresh Grace’s medium, and the time point was designated 0 h p.i. Cell supernatant was harvested at 12, 24, 48, 72, 96 and 120 h p.i., and viral titers were also determined for virus growth curve analysis (46). The viral titers were compared using F tests at each time point.

### Quantitative analysis of viral DNA synthesis and viral gene transcription

The qPCR analysis was performed as previously described (33) with some modifications. The RT-qPCR analysis was performed as previously described (47) with some modifications.

### Fluorescence confocal microscopy analysis

Confocal immunofluorescence microscopy was performed as previously described (48). Sf9 cells were seeded (1×10^6^ cells/dish) on a glass dish and allowed to attach for 2 h, then infected with vAc^*ac13Flag*REP^-*ph* at an MOI of 5. The cells were fixed with 4% paraformaldehyde for 10 min at the designated times. After washing with phosphate-buffered saline (PBS), the fixed cells were treated with 0.2% Triton X-100 for 10 min and then blocked with PBS containing 5% bovine serum albumin and 0.1% Tween-20 for 30 min. Subsequently, the cells were incubated with rabbit anti-FLAG polyclonal antibody (1:500, Proteintech) and mouse anti-Dm0 monoclonal antibody (1:500, DSHB) for 1 h followed by Alexa Fluor 594 goat anti-rabbit IgG (1:1000, Proteintech) and Alexa Fluor 647 goat anti-mouse IgG (1:1000, Proteintech) for 1 h in dark. Finally, the cells were stained with Hoechst 33258 (Beyotime, Shanghai, China) for 7 min in the dark and examined using a Leica SP5 confocal laser scanning microscope using a 60× dipping lens.

### TEM analysis

Sf9 cells were seeded (1×10^6^cells/dish) and allowed to adhere for 2 h, then infected with vAc^*ac13*KO^-*ph*, vAc^*ac13*REP^-*ph*, and vAc-*ph*. At 48 h p.i., the cells were fixed with 2.5% glutaraldehyde for 2 h for TEM as previously described (49). Ultrathin sections were visualized using a Tecnai G20 TWIN transmission electron microscope. Twenty intact cells infected with vAc^*ac13*KO^-*ph*, vAc^*ac13*REP^-*ph* or vAc-*ph* were randomly chosen to analyze nucleocapsid egress. The numbers of intranuclear and egressed nucleocapsids in each cell were counted using ImageJ software (https://imagej.nih.gov/ij/) and compared using the Kruskal-Wallis test followed by Dunn’s multiple comparison test.

### TEM, SEM and negative staining analysis

OBs were purified from larvae by differential centrifugation according to the method described by Gross et al (50) and observed by TEM (Hitachi Co., Ltd., Tokyo, Japan) and SEM (HITACHII SU-8010; Tokyo, Japan) as described previously (51). To observe ODVs embedded within OBs, 10 μL of OB suspension (10^8^ OBs/mL) were loaded onto a copper grid for 10 min. Filter paper was used to remove the remaining solution from the grid. Then, 10 μL of dissolution buffer was added to dissolve the OBs for 1 min. After removing the dissolution buffer, the grid was stained with 2% (w/v) phosphotungstic acid (pH 5.7) for 1 min. The grids were kept at room temperature overnight and observed by TEM. The ODVs in each OB were counted using ImageJ software (https://imagej.nih.gov/ij/) and their numbers were compared using the Kruskal-Wallis test followed by Dunn’s multiple comparison test.

## Acknowledgments

This work was supported by the National Key Research and Development Program of China (2017YFD0201206) and the WIV “One-Three-Five” strategic program (Y602111SA1 to XS). The funders had no role in study design, data collection and interpretation, or the decision to submit the work for publication. We thank the Core Facility and Technical Support of the Wuhan Institute of Virology, Chinese Academy of Sciences, for technical assistance with fluorescence microscopy (Ding Gao), flow cytometry (Juan Min), and electron microscopy (Pei Zhang, Anna Du and Bichao Xu).

## REFERENCES

1. Williams T, Bergoin M, van Oers MM. 2017. Diversity of large DNA viruses of invertebrates. J Invertebr Pathol 147:4–22.

2. Rohrmann GF. 2014. Baculovirus Molecular Biology (3rd edition). National Center for Biotechnology Information, Bethesda, MD.

3. Jehle JA, Blissard GW, Bonning BC, Cory JS, Herniou EA, Rohrmann GF, Theilmann DA, Thiem SM, Vlak JM. 2006. On the classification and nomenclature of baculoviruses: a proposal for revision. Arch Virol 151:1257–1266.

4. Herniou EA, Luque T, Chen X, Vlak JM, Winstanley D, Cory JS, O’Reilly DR. 2001. Use of whole genome sequence data to infer baculovirus phylogeny. J Virol 75:8117–8126.

5. Pearson MN, Rohrmann GF. 2002. Transfer, incorporation, and substitution of envelope fusion proteins among members of the Baculoviridae, Orthomyxoviridae, and Metaviridae (insect retrovirus) families. J Virol 76:5301–5304.

6. Slack J, Arif BM. 2006. The Baculoviruses Occlusion-Derived Virus: Virion Structure and Function, p. 99–165, Advances in Virus Research, vol. 69. Academic Press.

7. Fraser MJ. 1986. Ultrastructural Observations of Virion Maturation in Autographa-Californica Nuclear Polyhedrosis-Virus Infected Spodoptera-Frugiperda Cell-Cultures. J. Ultrastruct. Mol. Struct. Res. 95:189–195.

8. Young JC, MacKinnon EA, Faulkner P. 1993. The Architecture of the Virogenic Stroma in Isolated Nuclei of Spodoptera frugiperda Cells in Vitro Infected by Autographa californica Nuclear Polyhedrosis Virus. Journal of Structural Biology 110:141–153.

9. Johnson DC, Baines JD. 2011. Herpesviruses remodel host membranes for virus egress. Nat Rev Microbiol 9:382–394.

10. Hellberg T, Passvogel L, Schulz KS, Klupp BG, Mettenleiter TC. 2016. Nuclear Egress of Herpesviruses: The Prototypic Vesicular Nucleocytoplasmic Transport. Adv Virus Res 94:81–140.

11. Yue Q, Yu Q, Yang Q, Xu Y, Guo Y, Blissard GW, Li Z. 2018. Distinct Roles of Cellular ESCRT-I and ESCRT-III Proteins in Efficient Entry and Egress of Budded Virions of Autographa californica Multiple Nucleopolyhedrovirus. J Virol 92:e01636–01617.

12. Ohkawa T, Welch MD. 2018. Baculovirus Actin-Based Motility Drives Nuclear Envelope Disruption and Nuclear Egress. Curr Biol 28:2153–2159.

13. Guo Y, Yue Q, Gao J, Wang Z, Chen YR, Blissard GW, Liu TX, Li Z. 2017. Roles of Cellular NSF Protein in Entry and Nuclear Egress of Budded Virions of Autographa californica Multiple Nucleopolyhedrovirus. J Virol 91:e01111–01117.

14. Tao XY, Choi JY, Kim WJ, An SB, Liu Q, Kim SE, Lee SH, Kim JH, Woo SD, Jin BR, Je YH. 2015. Autographa californica multiple nucleopolyhedrovirus ORF11 is essential for budded-virus production and occlusion-derived-virus envelopment. J Virol 89:373–383.

15. Qiu J, Tang Z, Cai Y, Wu W, Yuan M, Yang K. 2019. The Autographa californica Multiple Nucleopolyhedrovirus *ac51* Gene Is Required for Efficient Nuclear Egress of Nucleocapsids and Is Essential for In Vivo Virulence. J Virol 93:e01923–01918.

16. Ke J, Wang J, Deng R, Wang X. 2008. Autographa californica multiple nucleopolyhedrovirus *ac66* is required for the efficient egress of nucleocapsids from the nucleus, general synthesis of preoccluded virions and occlusion body formation. Virology 374:421–431.

17. Guo YJ, Fu SH, Li LL. 2017. Autographa californica multiple nucleopolyhedrovirus *ac75* is required for egress of nucleocapsids from the nucleus and formation of de novo intranuclear membrane microvesicles. PLoS One 12:e0185630.

18. Shi A, Hu Z, Zuo Y, Wang Y, Wu W, Yuan M, Yang K. 2017. Autographa californica Multiple Nucleopolyhedrovirus *ac75* Is Required for the Nuclear Egress of Nucleocapsids and Intranuclear Microvesicle Formation. Journal of Virology 92:e01509–01517.

19. Tao XY, Choi JY, Kim WJ, Lee JH, Liu Q, Kim SE, An SB, Lee SH, Woo SD, Jin BR, Je YH. 2013. The Autographa californica multiple nucleopolyhedrovirus ORF78 is essential for budded virus production and general occlusion body formation. J Virol 87:8441–8450.

20. Olszewski J, Miller LK. 1997. A role for baculovirus GP41 in budded virus production. Virology 233:292–301.

21. Li Y, Shen S, Hu L, Deng F, Vlak JM, Hu Z, Wang H, Wang M. 2018. The functional oligomeric state of tegument protein GP41 is essential for baculovirus BV and ODV assembly. Journal of Virology 92:e02083–02017.

22. Yuan M, Huang Z, Wei D, Hu Z, Yang K, Pang Y. 2011. Identification of Autographa californica nucleopolyhedrovirus *ac93* as a core gene and its requirement for intranuclear microvesicle formation and nuclear egress of nucleocapsids. Journal of Virology 85:11664–11674.

23. Yuan M, Wu W, Liu C, Wang Y, Hu Z, Yang K, Pang Y. 2008. A highly conserved baculovirus gene *p48 (ac103*) is essential for BV production and ODV envelopment. Virology 379:87–96.

24. Fang M, Dai X, Theilmann DA. 2007. Autographa californica multiple nucleopolyhedrovirus EXON0 (ORF141) is required for efficient egress of nucleocapsids from the nucleus. Journal of Virology 81:9859–9869.

25. McCarthy CB, Dai X, Donly C, Theilmann DA. 2008. Autographa californica multiple nucleopolyhedrovirus *ac142*, a core gene that is essential for BV production and ODV envelopment. Virology 372:325–339.

26. Biswas S, Willis LG, Fang M, Nie Y, Theilmann DA. 2017. Autographa californica Nucleopolyhedrovirus AC141 (Exon0), a Potential E3 Ubiquitin Ligase, Interacts with Viral Ubiquitin and AC66 To Facilitate Nucleocapsid Egress. Journal of Virology 92:e01713–01717.

27. Chen YR, Zhong S, Fei Z, Hashimoto Y, Xiang JZ, Zhang S, Blissard GW. 2013. The transcriptome of the baculovirus Autographa californica multiple nucleopolyhedrovirus in Trichoplusia ni cells. J Virol 87:6391–6405.

28. Jones P, Binns D, Chang HY, Fraser M, Li W, McAnulla C, McWilliam H, Maslen J, Mitchell A, Nuka G, Pesseat S, Quinn AF, Sangrador-Vegas A, Scheremetjew M, Yong SY, Lopez R, Hunter S. 2014. InterProScan 5: genome-scale protein function classification. Bioinformatics 30:1236–1240.

29. Marchler-Bauer A, Bo Y, Han L, He J, Lanczycki CJ, Lu S, Chitsaz F, Derbyshire MK, Geer RC, Gonzales NR, Gwadz M, Hurwitz DI, Lu F, Marchler GH, Song JS, Thanki N, Wang Z, Yamashita RA, Zhang D, Zheng C, Geer LY, Bryant SH. 2017. CDD/SPARCLE: functional classification of proteins via subfamily domain architectures. Nucleic Acids Res 45:D200–D203.

30. Ono C, Kamagata T, Taka H, Sahara K, Asano S, Bando H. 2012. Phenotypic grouping of 141 BmNPVs lacking viral gene sequences. Virus Res 165:197–206.

31. Xiaoyong L, Keping C, Keya C, Qin Y. 2008. Determination of protein composition and host-derived proteins of Bombyx mori nucleopolyhedrovirus by 2-dimensional electrophoresis and mass spectrometry. Intervirology 51:369–376.

32. Kokusho R, Koh Y, Fujimoto M, Shimada T, Katsuma S. 2016. Bombyx mori nucleopolyhedrovirus BM5 protein regulates progeny virus production and viral gene expression. Virology 498:240–249.

33. Vanarsdall AL, Okano K, Rohrmann GF. 2005. Characterization of the replication of a baculovirus mutant lacking the DNA polymerase gene. Virology 331:175–180.

34. Shrestha A, Bao K, Chen YR, Chen W, Wang P, Fei Z, Blissard GW. 2018. Global Analysis of Baculovirus Autographa californica Multiple Nucleopolyhedrovirus Gene Expression in the Midgut of the Lepidopteran Host Trichoplusia ni. J Virol 92:e01277–01218.

35. Javed MA, Biswas S, Willis LG, Harris S, Pritchard C, van Oers MM, Donly BC, Erlandson MA, Hegedus DD, Theilmann DA. 2017. Autographa californica Multiple Nucleopolyhedrovirus AC83 is a Per Os Infectivity Factor (PIF) Protein Required for Occlusion-Derived Virus (ODV) and Budded Virus Nucleocapsid Assembly as well as Assembly of the PIF Complex in ODV Envelopes. J Virol 91:e02115–02116.

36. Huang Z, Pan M, Zhu S, Zhang H, Wu W, Yuan M, Yang K. 2017. The Autographa californica Multiple Nucleopolyhedrovirus *ac83* Gene Contains a cis-Acting Element That Is Essential for Nucleocapsid Assembly. J Virol 91:e02110–02116.

37. Victoria A, Mei Y, Carstens EB. 2009. Characterization of a baculovirus nuclear localization signal domain in the late expression factor 3 protein. Virology 385:209–217.

38. Feng G, Krell PJ. 2014. Autographa californica multiple nucleopolyhedrovirus DNA polymerase C terminus is required for nuclear localization and viral DNA replication. J Virol 88:10918–10933.

39. Olson VA, Wetter JA, Friesen PD. 2002. Baculovirus transregulator IE1 requires a dimeric nuclear localization element for nuclear import and promoter activation. J Virol 76:9505–9515.

40. Shen YW, Feng M, Wu XF. 2018. Bombyx mori nucleopolyhedrovirus ORF40 is essential for budded virus production and occlusion-derived virus envelopment. Journal of General Virology 99:837–850.

41. Klupp BG, Granzow H, Fuchs W, Keil GM, Finke S, Mettenleiter TC. 2007. Vesicle formation from the nuclear membrane is induced by coexpression of two conserved herpesvirus proteins. Proc Natl Acad Sci U S A 104:7241–7246.

42. Blissard GW, Theilmann DA. 2018. Baculovirus Entry and Egress from Insect Cells. Annu Rev Virol 5:113–139.

43. Datsenko KA, Wanner BL. 2000. One-step inactivation of chromosomal genes in Escherichia coli K-12 using PCR products. Proc Natl Acad Sci U S A 97:6640–6645.

44. Li J, Zhou Y, Lei C, Fang W, Sun X. 2015. Improvement in the UV resistance of baculoviruses by displaying nano-zinc oxide-binding peptides on the surfaces of their occlusion bodies. Appl Microbiol Biotechnol 99:6841–6853.

45. Li C, Wen A, Shen B, Lu J, Huang Y, Chang Y. 2011. FastCloning: a highly simplified, purification-free, sequence- and ligation-independent PCR cloning method. BMC Biotechnol 11:92.

46. Lei C, Yang J, Hu J, Sun X. 2020. On the Calculation of TCID50 for Quantitation of Virus Infectivity. Virol Sin.

47. Peng Y, Li K, Pei RJ, Wu CC, Liang CY, Wang Y, Chen XW. 2012. The protamine-like DNA-binding protein P6.9 epigenetically up-regulates Autographa californica multiple nucleopolyhedrovirus gene transcription in the late infection phase. Virol Sin 27:57–68.

48. Xu C, Wang J, Yang J, Lei C, Hu J, Sun X. 2019. NSP2 forms viroplasms during Dendrolimus punctatus cypovirus infection. Virology 533:68–76.

49. Qin F, Xu C, Hu J, Lei C, Zheng Z, Peng K, Wang H, Sun X. 2019. Dissecting the Cell Entry Pathway of Baculovirus by Single-Particle Tracking and Quantitative Electron Microscopic Analysis. J Virol 93:e00033–00019.

50. Gross CH, Russell RL, Rohrmann GF. 1994. Orgyia pseudotsugata baculovirus p10 and polyhedron envelope protein genes: analysis of their relative expression levels and role in polyhedron structure. J Gen Virol 75:1115–1123.

51. Kuang W, Zhang H, Wang M, Zhou NY, Deng F, Wang H, Gong P, Hu Z. 2017. Three Conserved Regions in Baculovirus Sulfhydryl Oxidase P33 Are Critical for Enzymatic Activity and Function. J Virol 91:e01158–01117.

